# Human skeletal muscle possesses an epigenetic memory of high intensity interval training

**DOI:** 10.1101/2024.05.30.596458

**Authors:** AM Pilotto, DC Turner, E Crea, R Mazzolari, L Brocca, MA Pellegrino, D Miotti, R Bottinelli, AP Sharples, S Porcelli

## Abstract

**INTRODUCTION:** Human skeletal muscle displays an epigenetic memory of resistance exercise induced by hypertrophy. It is unknown, however, whether high-intensity interval training (HIIT) also evokes an epigenetic muscle memory. This study employed repeated training intervention interspersed with a detraining period to assess epigenetic memory of HIIT.

**METHODS:** Twenty healthy subjects (25±5yrs) completed two HIIT interventions (training and retraining) lasting 2 months, separated by 3 months of detraining. Measurements at baseline, after training, detraining and retraining included maximal oxygen consumption (V̇O_2max_). Vastus lateralis biopsies were taken for genome-wide DNA methylation and targeted gene expression analyses. RESULTS: V̇O_2max_ improved during training and retraining (p<0.001) without differences between interventions (p>0.58). Thousands of differentially methylated positions (DMPs) predominantly demonstrated a hypomethylated state after training, retained even after 3-months exercise cessation and into retraining. Five genes; ADAM19, INPP5a, MTHFD1L, CAPN2, SLC16A3 possessed differentially methylated regions (DMRs) with retained hypomethylated memory profiles and increased gene expression. The retained hypomethylation during detraining was associated with an enhancement in expression of the same genes even after 3 months of detraining. SLC16A3, INPP5a, CAPN2 are involved in lactate transport and calcium signaling.

**CONCLUSIONS:** Despite similar physiological adaptations between training and retraining, memory profiles were found at epigenetic and gene expression level, characterized by retained hypomethylation and increased gene expression after training into long-term detraining and retraining. These genes were associated with calcium signaling and lactate transport. Whilst significant memory was not observed in physiological parameters, our novel findings indicate that human skeletal muscle possesses an epigenetic memory of HIIT.

## INTRODUCTION

Skeletal muscle memory refers to an enhanced response to a specific stimulus when the same or similar stimuli has been previously encountered (extensively reviewed in Sharples et al., 2016a; Sharples & Turner, 2023). In the context of exercise, muscle memory has wide implications for performance in recreational or elite athletes, or to inform potential strategies to maintain adaptation overtime in the general population to enable life-long muscle maintenance and therefore health and wellbeing (Sharples et al., 2015; Sharples et al., 2016b; Egan & Sharples, 2023). To date, muscle memory studies have primarily focused on adaptation to high-load resistance training where the muscle hypertrophic response is enhanced when the same resistance training stimulus has previously been applied (Staron et al., 1991).

To date, a positive muscle memory from exercise stimuli has been attributed to two mechanisms, ‘cellular’ memory and ‘epigenetic’ memory (epi-memory) that could also work in synergy (Sharples & Turner, 2023). Cellular memory suggests that skeletal muscle fibers retain increase in myonuclei accrued following an initial period of hypertrophy even when the anabolic stimulus ceases and muscle mass returns to baseline levels. The latter phenomenon would enable a favorable environment for more efficient and perhaps enhanced transcriptional and translational processes when muscle growth stimuli occur in the future. Such possibility was examined extensively in animal muscle using various mechanical loading/exercise mimicking regimes or steroid administration (Brusgaard et al., 2010; Egner et al., 2013; Gundersen, 2016; Lee et al., 2018; Murach et al., 2020; Eftestøl et al., 2022; Sharples & Turner, 2023). Recent meta-analysis of such studies suggested that the majority, yet perhaps not all myonuclei, are retained following such muscle growth stimuli (Rahmati et al., 2022). Whilst it is plausible that myonuclear retention after a physiologically induced training stimulus such as resistance training occurs in humans, evidence is not conclusive due to relatively few studies as well as to heterogeneity in the response to myonuclear accrual in humans (Psilander et al., 2019; Snijders et al., 2020; Blocquiaux et al., 2022). Indeed, muscle memory by myonuclear retention in humans is currently hotly debated (Murach et al., 2019; Eftestøl et al., 2020; Sharples & Turner, 2023).

As well as a positive memory by myonuclear retention, memory of exercise induced hypertrophy has been associated with retention of epigenetic modifications to the DNA in the skeletal muscle of humans and rodents. Seaborne and colleagues (Seaborne et al., 2018a), were the first to demonstrate that following resistance training, detraining and retraining in young adult humans, retraining evoked a larger increase in lower limb lean mass compared to the increment that occurred following the earlier training stimulus, even after muscle returned to pre-training mass during the intervening detraining period. The latter phenomenon was accompanied by epigenetic modifications (via DNA methylation) that revealed temporal profiles characterized by a predominance of retained DNA hypomethylation following the initial training period, even during detraining period, and/or followed by enhanced hypomethylation after later retraining (Seaborne et al., 2018a; Turner et al., 2019). Further, some memory gene signatures were associated with enhanced gene expression during retraining, which was also associated with enhancements in lean mass. Some of the epigenetic memory genes identified were subsequently demonstrated to be important in muscle mass regulation. For example, UBR5 was confirmed to be associated with hypertrophy and recovery from atrophy (Seaborne et al., 2019) and its knockdown evoked atrophy (Hughes et al., 2021). Since such initial evidence, Blocquiaux and colleagues (2022) also confirmed an epigenetic memory of resistance exercise-induced muscle growth in both young and aged adult human muscle (Blocquiaux et al., 2022). Wen and colleagues (Wen et al., 2021) also confirmed epigenetic muscle memory in mice using a model of training and detraining via progressive weighted wheel running that evokes hypertrophy. They also reported retained DNA methylation signatures into the detraining period following an earlier training period. Despite the progressive current research regarding muscle memory of muscle growth-induced exercise, there is currently little evidence for muscle memory of adaptation in response to divergent types of exercise including moderate continuous endurance and/or high intensity interval training that evoke alternate adaptations such as improved substrate utilization, oxidative capacity, and fatigue resistance.

To the authors knowledge, only a single study (Lindholm et al., 2016) investigated potential skeletal muscle memory at the transcriptional level in response to repeated endurance training interventions. Authors examined the transcriptome response to subsequent 12-week one-legged aerobic training periods (45 min, 4 times/week) interspersed by 9-month detraining. Results showed neither retention of gene expression profiles during the detraining period nor enhanced functional endurance adaptations during later retraining, which was not supportive of skeletal muscle memory following endurance training (Lindholm et al., 2016), at least at the transcriptional level. However, epigenetic modifications were not investigated. Recent studies in mice suggest enhanced and advantageous molecular response to acute exercise in already endurance trained muscle compared with naive/untrained muscle due to a ‘priming’ of an advantageous epigenetic landscape via DNA methylation (Furrer et al., 2023). Despite this, epigenetic memory of endurance or high intensity exercise training has not been studied in humans using a training, detraining, retraining model yet. To further support the rationale for this study, it has also been previously demonstrated that acute aerobic exercise at higher exercise intensities (80% V̇O_2max_) compared with lower intensities (40% V̇O_2max_) evokes hypomethylation of PGC-1α pathway and mitochondrial biogenesis target genes (Barrès et al., 2012). Moreover, across the genome, we have previously demonstrated that acute sprint interval exercise at higher intensities evokes greater hypomethylation of genes in exercise related pathways such as the AMPK, MAPK pathway and axon guidance pathways (Maasar et al., 2021). Among the latter genes, several play a major role in exercise adaptation to oxidative metabolism (e.g. NR4A1 & 3), and capillarization (e.g. VEGFA) (Pillon et al., 2020; Maasar et al., 2021). Therefore overall, together with this information and that retained hypomethylation of muscle growth-related genes was observed after muscle growth in humans (Seaborne et al., 2018a; Turner et al., 2019; Blocquiaux et al., 2022), it is hypothesized that high intensity interval training may also evoke an epigenetic memory (via retained hypomethylation), albeit perhaps in metabolic and mitochondrial-related genes.

The aim of the present study was to investigate whether human skeletal muscle possesses an epigenetic memory of repeated high intensity interval training (HIIT) separated by a long-term period of detraining (3 months) by examining the response at genome wide DNA methylome level and identifying key epigenetic memory genes and pathways.

## MATERIALS AND METHODS

### Subjects

The study intervention was conducted at the University of Pavia, Italy. Twenty male (n = 11) and female (n = 9) adults were recruited from the local community. Participants anthropometrics and physiological characteristics are reported in **Table 1**. Inclusion criteria included: To have not been involved in structured training programs before; maximal oxygen consumption (V̇O_2max_), determined by an initial screening test, lower than 40 and 45 ml·min^−1^·kg^−1^ for women and men, respectively. Exclusion criteria were assessed by both questionnaires and physical screening to identify that all participants were free from major diseases at the pulmonary, cardiovascular and muscle level and they were not taking any drugs. All procedures were in accordance with the Declaration of Helsinki and the study was approved by the local ethics committee (Besta 64-19/07/2019) and participants gave their written informed consent after being informed about objectives, methods and risks of the study.

**Table 1.** Participants anthropometrics and physiological characteristics measured during the initial screening test grouped according to sex. V̇O_2max_, maximal oxygen uptake. All values are means ± SD.

| | Age<br>(years) | Weight<br>(kg) | Height<br>(cm) | $\dot{V}O_{2max}$<br>(ml·min<sup>-1</sup>·kg<sup>-1</sup>) | $\dot{V}O_{2max}$<br>(l·min<sup>-1</sup>) |
| --- | --- | --- | --- | --- | --- |
| Males (n = 11) | 27 ± 5 | 72.1 ± 10.7 | 177 ± 5 | 36.8 ± 8.3 | 2.601 ± 0.448 |
| Females (n = 9) | 23 ± 3 | 58.7 ± 7.2 | 167 ± 4 | 34.1 ± 5.3 | 1.985 ± 0.308 |

### Study design

The current study exposed the same individuals to two identical 8-week (2 months) high intensity interval training periods separated by a three-month washout period during which participants were instructed to return to their habitual life (total approx. 7 months). Levels of physical activity were monitored before and throughout the entire period of the study by accelerometers (wGT3X-BT, ActiGraph, Florida, USA). Data collection was performed at four different time points: at baseline (BASELINE), after the first training period (TRAINING), following detraining (DETRAINING) and after retraining (RETRAINING). For each time point participants visited the laboratory on two separate occasions. During the first visit, anthropometric measurements were collected, and an incremental cycle ergometer exercise test up to voluntary exhaustion was performed. On the second visit, approximately 100 mg of skeletal muscle tissue was obtained from the *vastus lateralis* muscle under local anesthesia (1% lidocaine) for genome-wide DNA methylation (methylome), targeted gene expression based on genes that demonstrated significantly differentially methylation positions (DMPs) or regions (DMRs) (described below) demonstrating retained epigenetic profiles. Participants were instructed to abstain from strenuous physical activity for at least 48 h prior to each testing session. Experimental design is reported in **Figure 1**.

**Figure 1.**
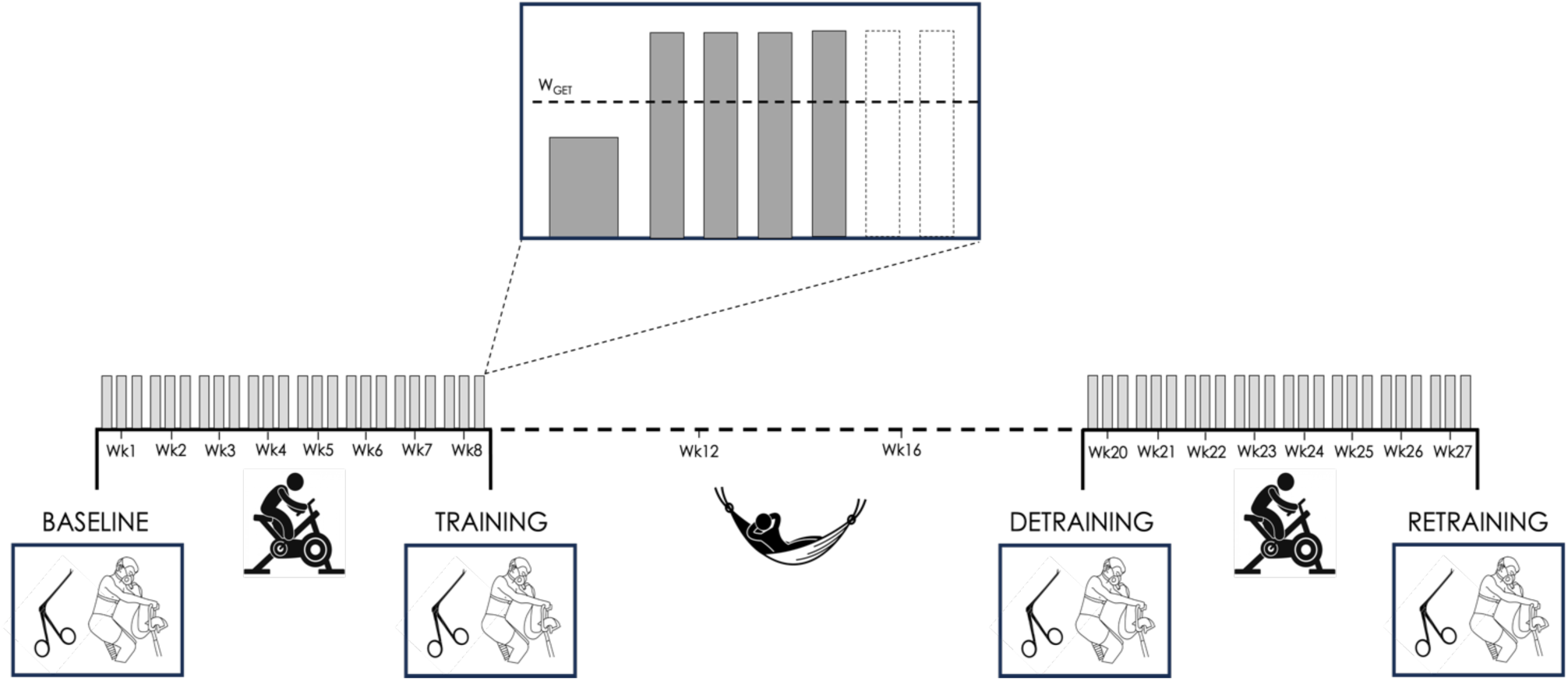
Experimental design. Individuals were exposed to two identical 8-week high intensity interval training periods separated by a 3-month washout period (detraining) during which participants were instructed to return to their habitual physical activities. Incremental maximal cycle ergometer exercise tests and muscle biopsies were performed at four different time points: at baseline (BASELINE), after the first training period (TRAINING), following detraining (DETRAINING) and after retraining (RETRAINING). W_GET_, power output at gas exchange threshold.

### Training

Exercise prescription consisted of two identical 8-week periods of high intensity interval training (HIIT) based on individual results obtained from the exercise incremental tests. The first three training sessions were conducted under the supervision of a researcher, who was in charge to give instructions to the participants about the training procedures. The remaining training sessions (3/week) were conducted at home on electronically braked cycle ergometer. Time, heart rate (HR) and power (W) from each session were recorded and uploaded on a web-based app (MZ3, MyZone, England, UK) to allow the researchers to check the adherence to the training regimen. Every week the subjects were contacted by phone to monitor progression, provide feedback and encouragement, and to answer any questions. Exercise sessions were composed by 10 min of warm-up followed by high intensity bouts of short (1-2 min) and long (3-4 min) duration. Exercise prescription varied during the training period to facilitate participant motivation and compliance, as detailed previously (Robach et al., 2014). To avoid stagnation (Granata et al., 2016) the training stimulus (power and repetitions) was progressively incremented. An identical training protocol was applied for both training and retraining.

### Incremental exercise

Incremental exercise was performed on an electronically braked cycle ergometer (818E, Monark, Sweden) to determine maximal oxygen consumption (V̇O_2max_), gas exchange threshold (GET), and respiratory compensation point (RCP). Pedaling frequency was digitally displayed to the subjects, and subjects were asked to keep a constant cadence throughout the tests between 70 and 80 rpm. Power was increased 10-15 W every minute, depending on the individual’s fitness aiming to allow the subjects to reach voluntary exhaustion in 10–15 min. Voluntary exhaustion was defined as the incapacity to maintain pedaling frequency for 5 s at the imposed work rate despite vigorous encouragement by the researchers. After 30 min of recovery, all participants performed a verification trial consisting of a constant work rate exercise at 90% of the highest power attained in the incremental test (W_max_) (Rossiter et al., 2006).

Pulmonary ventilation (V̇_E_, in BTPS [body temperature (37°C), ambient pressure and gas saturated with water vapor]), oxygen consumption (V̇O_2_), and CO_2_ output (V̇CO_2_), both in STPD (standard temperature [0°C or 273 K] and pressure [760 mmHg] and dry [no water vapor]), were determined breath-by-breath by a metabolic cart (Vyntus CPX, Vyaire Medical GmbH, Germany). Before each test, gas analyzers were calibrated with ambient air and a gas mixture of known concentration (O_2_: 16%, CO_2_: 4%) and the turbine flowmeter was calibrated with a 3-L syringe at three different flow rates. RER was calculated as V̇CO_2_/V̇O_2_. HR was recorded by using a HR chest band (HRM-Dual, Garmin, Kansas, USA). Rating of perceived exertion (RPE) was determined using Borg® 6–20 scale every 2 min through the test (Borg, 1982). At rest, and at 1, 3, and 5 min of recovery, 20 μL of capillary blood was obtained from a preheated earlobe for blood lactate concentration (Biosen C-line, EKF, Germany); the analyzer was frequently calibrated with a standard solution containing 12 mmol·L^−1^ of lactate. The highest 20-s averaged cardiopulmonary and metabolic data recorded during the whole incremental test (i.e. including the verification phase) were taken as maximal values (Martin-Rincon et al., 2019). GET was determined by two independent investigators by using the modified “V-slope” method (Beaver et al., 1986) and “secondary criteria”. RCP was determined as the point where end-tidal PCO_2_ began to fall after a period of isocapnic buffering (Whipp et al., 1989).

### Muscle Biopsy

Fifteen participants out of 20 [male (n = 9) and female (n = 6)] agreed to undergo muscle biopsy for all the time points required by the experimental protocol. Resting muscle biopsies were taken from the *vastus lateralis* muscle using a 130 mm (6”) Weil-Blakesley rongeur (NDB-2, Fehling Instruments, GmbH&Co, Germany) under local anesthesia (1% lidocaine). After collection, muscle samples were cleaned of excess blood, fat, and connective tissue and divided into different portions. A portion of ∼20 mg of muscle tissue was immediately submerged in All protect Tissue Reagent (Qiagen, Germany) following the manufacturer’s instructions to stabilize and protect cellular DNA and was stored at –20°C for subsequent DNA methylome analysis (described below). A specimen of ∼20-30 mg was rapidly frozen by immersion in liquid nitrogen and stored at –80°C for RNA isolation and gene expression analysis (described below). Fifteen out of 15 subjects were analyzed for RNA/gene expression, and 5 subjects per time point (BASELINE, TRAINING, DETRAINING, RETRAINING) for DNA methylome analysis.

### Methylation analysis

After power analysis conducted to detect a greater than 1.05 (5%) fold change in methylation based on our previous studies (Seaborne et al., 2018a), a sample size of 4 in a within-subject design was determined as sufficient to detect statistically significant changes in methylation over the baseline, training, detraining and retraining time points. Thereby, five (*n* = 5) subjects were analyzed in the present study for DNA methylome analysis.

### Tissue Homogenization, DNA Isolation, and Bisulfite Conversion

Tissue samples were homogenized for 45 s at 6,000 rpm × 3 (5 min on ice in between intervals) in lysis buffer (180 μl buffer ATL with 20 μl proteinase K) provided in the DNeasy spin column kit (Qiagen, Germany) using a Roche Magnalyser instrument and homogenization tubes containing ceramic beads (Roche, Germany). The DNA was then bisulfite converted using the EZ DNA Methylation Kit (Zymo Research, CA, USA) as per manufacturer’s instructions.

### Infinium Methylation EPIC Beadchip array

All DNA methylation experiments were performed in accordance with Illumina manufacturer instructions for the Infinium Methylation EPIC 850K BeadChip Array (Illumina, CA, USA). Methods for the amplification, fragmentation, precipitation and resuspension of amplified DNA, hybridization to EPIC BeadChip, extension and staining of the bisulfite converted DNA was conducted as detailed in paper from Seaborne and colleagues (Seaborne et al., 2018b). EPIC BeadChips were imaged using the Illumina iScan System (Illumina, CA, USA). DNA methylation analysis, CpG enrichment analysis (GO and KEGG pathways), differentially methylated region (DMR) analysis and Self Organizing Map (SOM) temporal profiling were performed. Following DNA methylation quantification via Methylation EPIC BeadChip array, raw.IDAT files were processed using Partek Genomics Suite V.7 (Partek Inc., Missouri, USA) and annotated using the MethylationEPIC_v-1-0_B4 manifest file. We first checked the average detection p-values for each sample across all probes. The mean detection p-value for all samples across all probes was 0.000295, and the highest for any given sample was 0.000597, which is well below the recommended 0.01 in the Oshlack workflow (Maksimovic et al., 2016). We also produced density plots of the raw intensities/signals of the probes per sample. These demonstrated that all methylated and unmethylated signals were over 11.5 (mean median signal for methylated probes was 11.56 and unmethylated probes 11.69), and the mean difference between the median methylation and median unmethylated signal was 0.13, well below the recommended difference of less than 0.5 (Maksimovic et al., 2016). Upon import of the data into Partek Genomics Suite we removed probes that spanned X and Y chromosomes from the analysis due to having both males and females in the study design, and although the average detection p-value for each sample was very low (no higher than 0.000597) we also excluded any individual probes with a detection p-value that was above 0.01 as recommended (Maksimovic et al., 2016). Out of a total of 865,859 probes removing those on the X & Y chromosome (19,627 probes) and with a detection p-value above 0.01 (4,264 probes) reduced the total probe number to 843,355 (note some X&Y probes also had detection p-values of above 0.01). We also filtered out probes associated with single-nucleotide polymorphisms (SNPs) and any known cross-reactive probes using previously defined SNP and cross-reactive probe lists from EPIC BeadChip 850K validation studies (Pidsley et al., 2016). This resulted in a final list of 791,084 probes to be analysed. Following this, background normalization was performed via functional normalization (with noob background correction) as previously described (Maksimovic et al., 2012). Following functional normalization, we also undertook quality control procedures via Principle Component Analysis (PCA), density plots by lines as well as box and whisker plots of the normalized data for all samples. Any outlier samples were detected using Principle Component Analysis (PCA) and the normal distribution of β-values. Outliers were detected if they fell outside 2 standard deviations (SDs) of the ellipsoids and/or if they demonstrated different distribution patterns to the samples of the same condition. We confirmed that no samples demonstrated large variation [variation defined as any sample above 2 standard deviations (SDs) – depicted by ellipsoids in the PCA plots and/or demonstrating any differential distribution to other samples, depicted in the signal frequency by lines plots. Therefore, no outliers were detected in this sample set. Following normalization and quality control procedures, we undertook differentially methylated position (DMP) analysis by converting β-values to M-values [M-value = log2(β/(1 − ββ)], as *M*-values show distributions that are more statistically valid for the differential analysis of methylation levels (Du et al., 2010). We undertook a one-way ANOVA for comparisons of baseline, training, detraining and retraining. Any differentially methylated CpG position (DMP) with an unadjusted p-value of ≤ 0.01 was used as the statistical cut off for the discovery of DMPs. We then undertook CpG enrichment analysis on these differentially methylated CpG lists within gene ontology (GO) and KEGG pathways (Kanehisa & Goto, 2000; Kanehisa et al., 2016; Kanehisa et al., 2017) using Partek Genomics Suite and Partek Pathway. Differentially methylated region (DMR) analysis, that identifies where several CpGs are consistently differentially methylated within a short chromosomal location/region, was undertaken using the Bioconductor package DMRcate (DOI: 10.18129/B9.bioc.DMRcate). Finally, to plot and visualize temporal changes in methylation across the timepoints we implemented Self Organizing Map (SOM) profiling of the change in m-value within each condition using Partek Genomics Suite.

### Gene Expression Analysis

Gene Expression Analysis was performed on muscle samples from fifteen (n = 15) subjects.

### Tissue Homogenization, RNA extraction and RT-PCR

Skeletal muscle tissue muscle was homogenized in tubes containing ceramic beads (MagNA Lyser Green Beads, Roche, Germany) and 1 ml Tri-Reagent (Invitrogen, England, UK) for 45 s at 6,000 rpm×3 (and placed on ice for 5 min at the end of each 45 s homogenization) using a Roche MagNA Lyser instrument (Roche, Germany). RNA was extracted using standard Tri-Reagent procedure via chloroform/isopropanol extractions and 75% ethanol washing as per manufacturer’s instructions. RNA pellets were resuspended in RNA storage solution (Ambion, England, UK) and analyzed (Nanodrop, ThermoFisher Scientific, England, UK) for an indication of quality (260/280 ratio of mean ± SD, 2.01 ± 0.10). Then a one-step RT-PCR reaction (reverse transcription and PCR) was performed using QuantiFast SYBR® Green RT-PCR one-step assay kits on a Rotorgene 3000Q. Each reaction was setup as follows; 4.75 μl experimental sample (7.36 ng/μl totalling 35 ng per reaction), 0.075 μl of both forward and reverse primer of the gene of interest (100 μM stock suspension), 0.1 μl of QuantiFast RT Mix (Qiagen, Germany) and 5 μl of QuantiFast SYBR Green RT-PCR Master Mix (Qiagen, Germany). Reverse transcription was initiated with a hold at 50 °C for 10 min (cDNA synthesis), followed by a 5-min hold at 95 °C (transcriptase inactivation and initial denaturation), before 40–50 PCR cycles of; 95 °C for 10 s (denaturation) followed by 60 °C for 30 s (annealing and extension). Primer sequences for genes of interest and reference genes are included in **Table 2**. Primers were designed to detect a single target gene and, if application, its associated transcript variants. All genes demonstrated no unintended targets via BLAST search and yielded a single peak after melt curve analysis conducted after the PCR step above. All relative gene expression was quantified using the comparative Ct (ΔΔCt) method. Individual participants own baseline Ct values were used in ΔΔCt equation as the calibrator using RPL13a as the reference gene. The average Ct value for the reference gene was consistent across all participants and experimental conditions (18.57 ± 0.81, SDEV) with low variation of 4.36%.

**Table 2.**
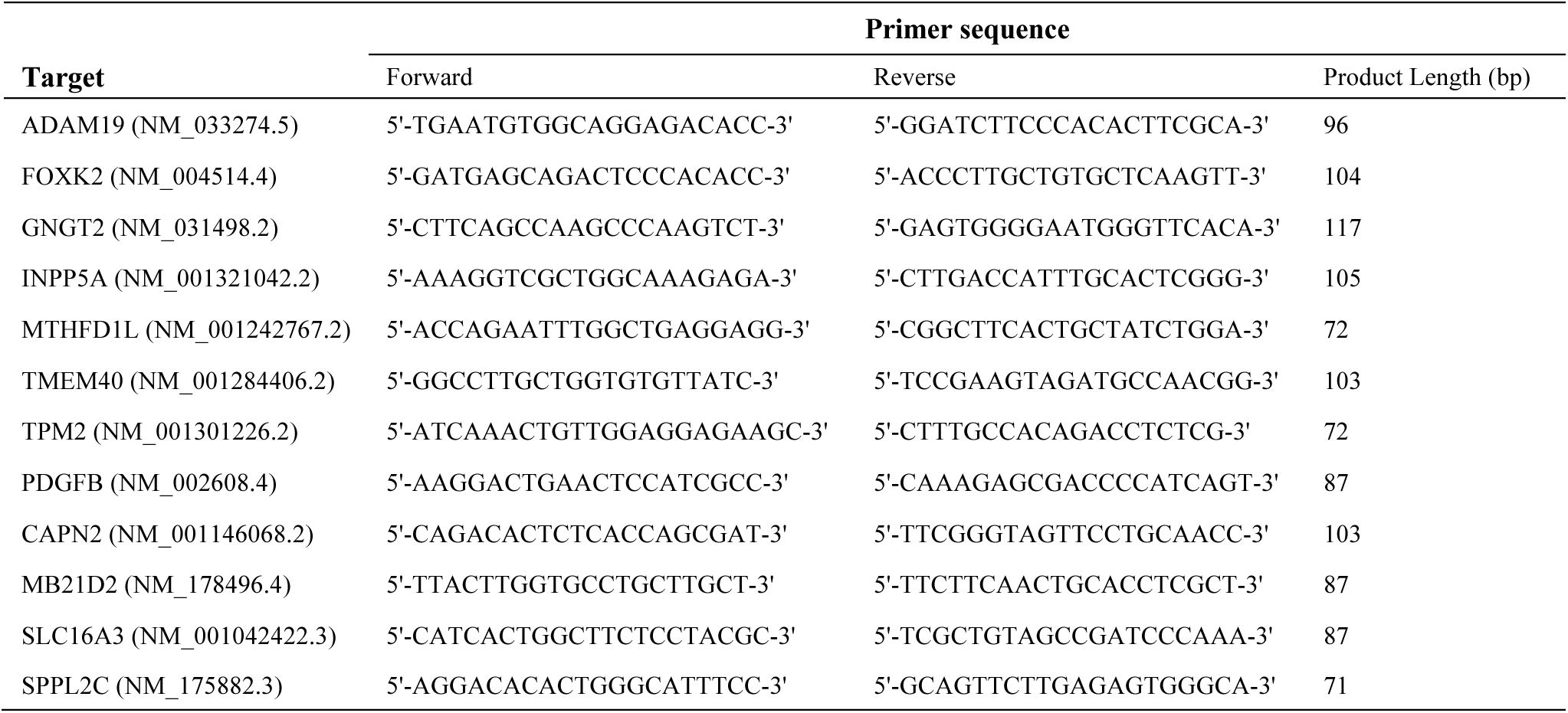
Primer sequences for real-time PCR.

### Statistical Analysis

Results are reported as the mean ± SD unless otherwise stated. Outcome measures were assessed using one-way repeated measure ANOVA to determine significant overall main effects of time (BASELINE, TRAINING, DETRAINIG, RETRAINING). Once an overall effect was confirmed, statistical significance of the measured difference among groups was assessed by Tukey’s post hoc analysis. Statistical values were considered significant at the level of p ≤ 0.05, and DNA methylation analysis where statistical significance was placed at a p ≤ 0.01. Adjusted p values with the corresponding 95% confidence interval are reported. For epigenetics analysis methylome wide array data sets for baseline, training and retraining were analyzed for significant DMPs in Partek Genomics Suite. All gene ontology and KEGG signaling pathway analysis was performed in Partek Genomics Suite and Partek Pathway, on generated DMPs list across conditions (ANOVA) or contrasts between paired conditions. For follow up M-value difference in CpG DNA methylation analysis was performed via ANOVA in MiniTab Statistical Software (MiniTab Version 17.2.1). Follow up RT-qRT-PCR gene expression was analyzed using both a repeated measures one-way ANOVA, to detect for significant changes across time for identified clusters of genes, and an ANOVA for follow up of individual genes over time.

## RESULTS

### Physiological effects of training

The average intensity prescribed across all 24 training sessions for each training period was set to be 120% of individual W_max_ relative to baseline for the initial training intervention and in relation to detraining for the retraining phase. The absolute workload effectively carried out by participants in the initial training period was 245 ± 52 W, and this did not exhibit any statistical difference from the power output recorded during the subsequent retraining period (247 ± 53 W, p = 0.653) (**Fig. 2A**). Subjects achieved an average intensity of 202% and 208% of W_GET_ during training and retraining, respectively (p = 0.504) (**Fig. 2B**).

**Figure 2.**
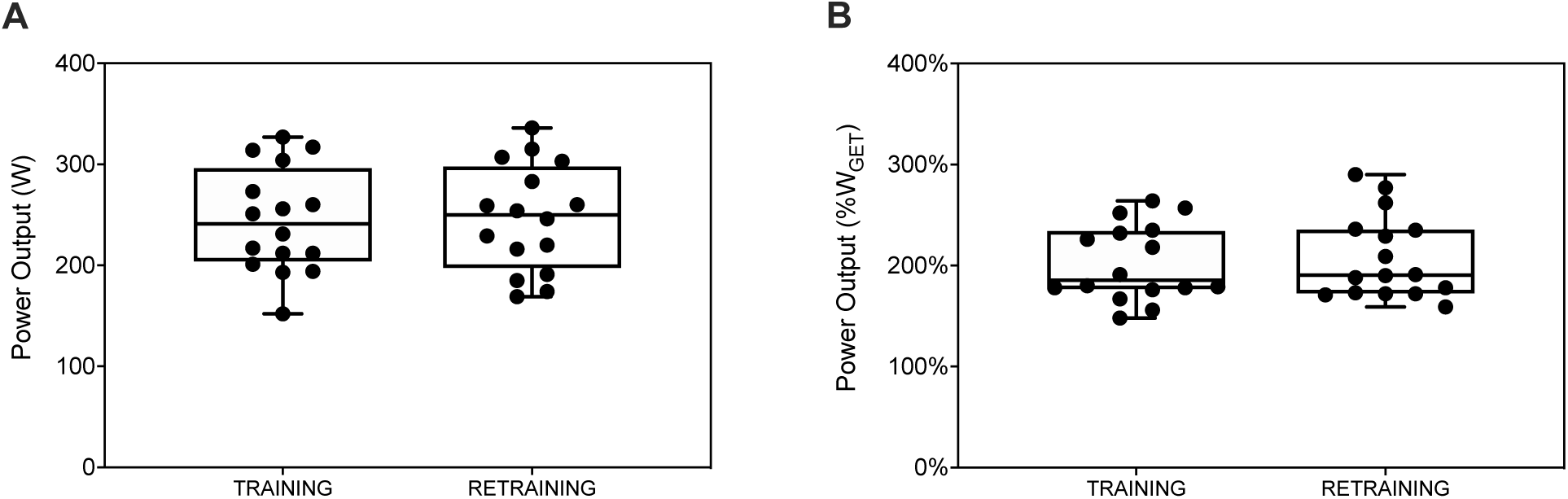
Average training intensity performed by individuals during training and retraining. Workload effectively performed by participants in absolute values (panel A) and in relation to W_GET_ assessed during the incremental exercise conducted before each training period (panel B) (n = 20). W_GET_, power output at gas exchange threshold.

Maximal values of the main respiratory, cardiovascular, and metabolic variables obtained during the incremental exercise are shown in **Table 3**. W_max_ and V̇O_2max_ increased significantly after TRAINING (+18% and +14%, respectively; both p < 0.001) and RETRAINING (+13% and +14%, respectively; both p < 0.001). DETRAINING reported maximal values of all the variables back to BASELINE levels (all p > 0.99). HR_max_ was between 87 and 113% of the age-predicted maximum values [calculated as 208 ± 0.7 · age (Tanaka et al., 2001)]. RER_max_, [La]_b max_, and RPE_max_ confirmed that subjects exercised until the limit of tolerance, resulting in no significant difference amongst the time-points.

**Table 3.**
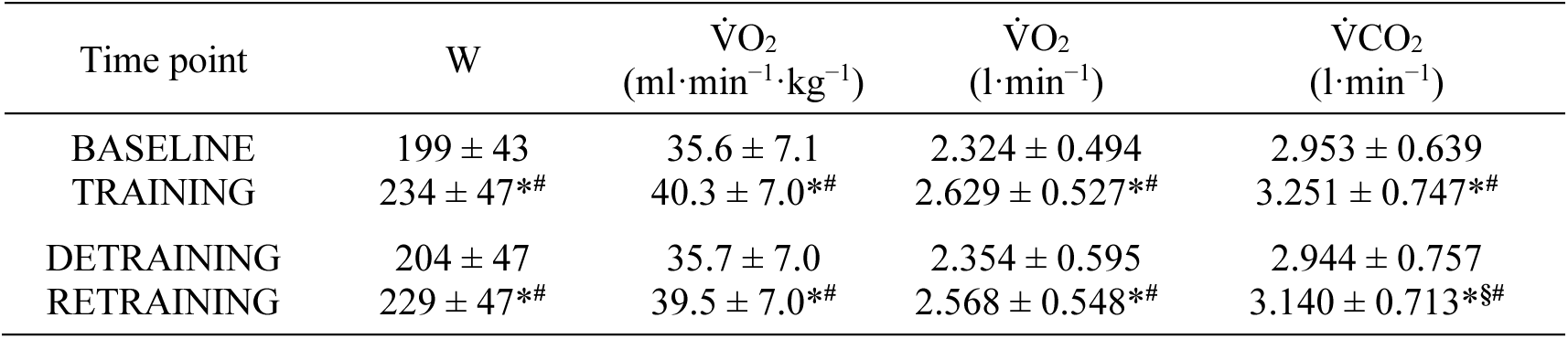
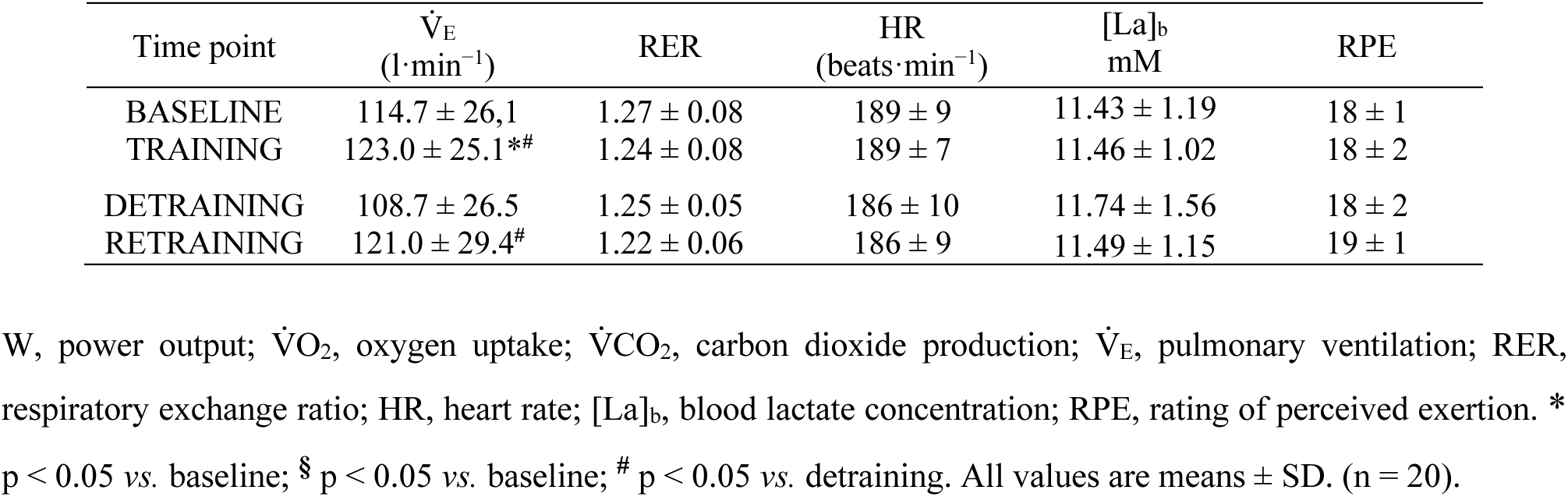
Maximal values of main respiratory, cardiovascular, and metabolic variables determined during incremental exercise.

### Genome-wide DNA methylation

The frequency of differentially methylated CpG positions (DMPs) in each condition was analyzed (**Fig. 3**) and a list of 35,855 DMPs were identified. Compared to BASELINE, 21,605 CpG sites were significantly differentially methylated following TRAINING with a larger number being hypomethylated (n=14,516) compared to hypermethylated (n=7,089). After DETRAINING, the total number of DMPs decreased (n=3,854) but did not revert back to BASELINE levels. Most of them were hypomethylated (n=3,349), whereas hypermethylated DNA sites nearly reverted to BASELINE (n=505). Following RETRAINING, an increase in the number of DMPs was observed (n=10,396) compared to DETRAINING and BASELINE, with most DMPs hypomethylated (n=9,644). Indeed, the number of hypermethylated DNA sites remained similar to DETRAINING (n=752). The increment in hypomethylated DMPs during RETRAINING was smaller compared to TRAINING (a list of all the identified DMPs for all comparisons can be downloaded in **Suppl. File S1**). Interestingly, the larger increase in hypomethylation was associated with a much smaller number of DMPs that were hypermethylated after RETRAINING compared to TRAINING (n=752 vs. n=7,089). Further, when DMPs are expressed as a percentage of total differentially methylated sites from BASELINE, there was a positive trend towards an enhanced proportion of hypomethylated positions across time-points (67%, 87% and 93% in TRAINING, DETRAINING and RETRAINING, respectively) (**Fig. 3B**). Regarding genic region-specific methylation, 64%, 45% and 90% of promoter associated DMPs were hypomethylated following TRAINING, DETRAINING and RETRAINING, respectively, resulting in nearly 30% more promoter-associated hypomethylated DMPs after RETRAINING (90%) versus TRAINING (64%) (**Suppl. File S1**). The positions of all DMPs in relation to CpG islands and gene regulatory groups can be found in **Suppl. File S2**. Gene Ontology (GO) and KEGG pathway analyses indicated that differential methylation that occurred at all timepoints of TRAINING, DETRAINING and RETRAINING was enriched in 8 common pathways across time of: (1) axon guidance, (2) calcium signaling pathway, (3) cholinergic synapse, (4) circadian entrainment, (5) focal adhesion, (6) MAPK signaling pathway, (7) pathways in cancer, (8) Rap1 signaling pathway (all comparisons and overlapping common pathways are in **Suppl. File S3** and **Suppl. File S4**, respectively).

**Figure 3.**
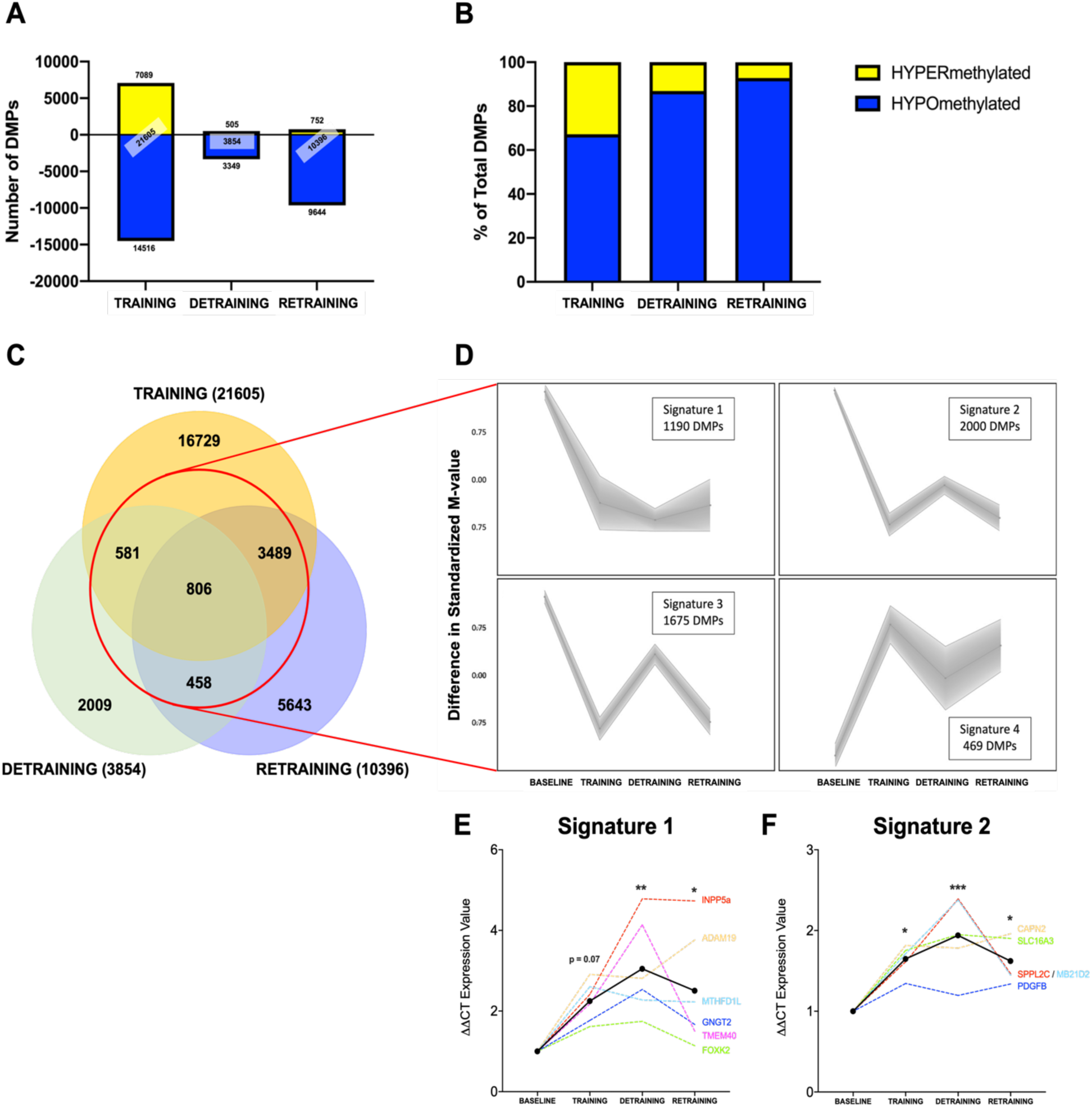
**A**) Total number of DMPs hypo-(blue) and hyper-methylated (yellow) following TRAINING, DETRAINING and RETRAINING conditions compared to baseline (n = 5). **B**) Hypo-(blue) and hypermethylated (yellow) DMPs as a percentage of the total number of DMPs (n = 5). **C**) Venn diagram analysis of the statistically differentially regulated CpG sites following TRAINING, DETRAINING and RETRAINING compared to BASELINE. Ellipses report the number of common overlapping significant DMPs across each condition (n = 5). **D**) SOM profiling depicting the temporal regulation of DMPs over the time-course of BASELINE, TRAINING, DETRAINING and RETRAINING. Signature 1 DMPs were hypomethylated after TRAINING and maintained hypomethylation even after 3 months DETRAINING and RETRAINING. Signature 2 DMPs were hypomethylated after TRAINING; hypomethylated DMPs were reduced slightly during DETRAINING yet still retained as hypomethylated relative to BASELINE, then after RETRAINING demonstrated similar or in some cases (see main text) an enhanced hypomethylated state compared with TRAINING. Signature 3 DMPs were hypomethylated after TRAINING, returning towards BASELINE levels after DETRAINING and then hypomethylated once again after RETRAINING. Signature 4 DMPs were hypermethylated after TRAINING, returning slightly towards BASELINE levels and then hypermethylated once more after RETRAINING. **E**) Gene expression determined by RT-PCR for the cluster of genes identified in signature 1 (ADAM19, FOXK2, GNGT2, INPP5a, MTHFD1L, TMEM40) showing an increase in gene expression after TRAINING that continued to increase despite training completely ceasing during DETRAINING and was retained as upregulated after RETRAINING. * p < 0.05, ** p < 0.01 compared to baseline. All significance taken as p less than or equal to 0.05 unless otherwise reported on graph (n = 15). **F**) Expression of genes identified in signature 2 (CAPN2, PDGFB, MB21D2, SLC16A3, SPPL2C) showing a significant increase in expression after TRAINING continuing even during DETRAINING and remaining elevated after RETRAINING. * p < 0.05, *** p < 0.001 compared to baseline (n = 15).

In order to investigate whether there were any similarly altered DMPs across interventions, we overlapped DMP lists from analysis described above to detect differentially methylated CpG sites in the three conditions versus BASELINE. Around 15% DMPs (n=5,334) overlapped and 806 DMPs were modified in all time-points (**Fig. 3C**). Changes in genome-wide DNA methylation related to the overlapped DMPs (n=5,334) were further analyzed by self-organizing map (SOM) profiling (or clustering). SOM highlighted four temporal trend profiles (**Fig. 3D**). The first temporal profile (signature 1) included DMPs (n=1,190) that were hypomethylated after TRAINING and retained hypomethylation even after 3 months of DETRAINING and subsequent RETRAINING. The second temporal profile (signature 2) included DMPs (n=2,000) that were also hypomethylated after TRAINING. Hypomethylation was only slightly reduced relative to BASELINE during DETRAINING and went back to training levels after RETRAINING. The third profile (signature 3) included DMPs (n=1,675) which were also hypomethylated after TRAINING but returned towards BASELINE levels after DETRAINING and were hypomethylated once again after RETRAINING. The final profile (signature 4) included DMPs (n=469) that were hypermethylated after TRAINING, returned slightly towards BASELINE levels after DETRAINING and then hypermethylated once again after RETRAINING. Overall, such temporal profiling indicated that the majority of DMPs were hypomethylated with training (signature 1-3) and that an epigenetic memory of retained hypomethylation after detraining and retraining (signature 1 & 2) occurred. Retention of hypermethylation was also detected (signature 4), albeit in a smaller number of DMPs (n=469) compared with a combined total of 3,190 DMPs with retained hypomethylation (signature 1 & 2). Interestingly, the Wnt pathway, involved in activation of growth-control genes in skeletal muscle (Armstrong & Esser, 2005), was the top enriched pathway in memory signature 1 that showed the most prominent retained hypomethylation from the earlier training into detraining and retraining. All DMPs identified in each signature 1-4 can be accessed in **Suppl. Files S5** and their gene ontology and KEGG pathway analysis in **Suppl. Files S6**.

### Gene Expression – Identification of epigenetic memory gene profiles and their gene expression

According to the analysis of the DMPs identified in the epigenetic memory signatures above, we first determined which of the genes demonstrated multiple DMPs within short chromosomal regions of the same gene, also known as differentially methylated regions (DMRs). This is because differential methylation in multiple places of the same region is perhaps more likely to functionally impact gene transcription. DMRs with a signature 1 profile were identified across 6 genes: *ADAM19, FOXK2, GNGT2, INPP5a, MTHFD1L, TMEM40*. Five DMRs were identified with the signature 2 profile: *CAPN2, MB21D2, SLC16A3, SPPL2C, AL031590.1 (which is antisense for the PDGFB gene)*. For signature 3, no genes with DMRs were detected, and Signature 4 contained one gene that contained a DMR (TPM2). All DMRs identified in each signature can be accessed in **Suppl. File S7**. We then ran targeted gene expression using RT-qPCR on such memory profile genes. The 6 genes identified by DMRs in signature 1 gene cluster (ADAM19, FOXK2, GNGT2, INPP5a, MTHFD1L, and TMEM40) showed a significant main effect for time (p < 0.01) after repeated measures and one-way ANOVA analysis. After 8 weeks of high intensity interval training, this cluster displayed non-significant increased gene expression (2.25 ± 0.50, p = 0.07) (**Fig. 3E**). Importantly, even when exercise TRAINING was completely ceased for 3 months (DETRAINNG) gene expression was enhanced (3.05 ± 1.17) resulting in statistical significance (p < 0.01) compared to BASELINE. Enhanced gene expression was then retained (2.50 ± 1.43) after RETRAINING compared to BASELINE (p < 0.05) (**Fig. 3E**). Repeated measures and one-way ANOVA analysis for the 5 genes identified by DMRs in signature 2 cluster (CAPN2, MB21D2, SLC16A3, SPPL2C and PDGFB) also reported a significant effect of time (p < 0.01). Upon TRAINING, gene expression of this cluster significantly increased (1.65 ± 0.18, p < 0.05) and showed further increase even after DETRAINING (1.94 ± 0.49, p < 0.001) and RETRAINING (1.62 ± 0.29, p < 0.05) compared to BASELINE (**Fig. 3F**). Closer analysis of CpG DNA methylation of the 11 gene clusters, identified a distinct inverse relationship between methylation and gene expression in 5 gene: ADAM19, INPP5a, MTHFD1L, CAPN2 and SLC16A3 (**Fig. 4**). Upon 8 weeks of high intensity interval TRAINING, hypomethylation of such genes was associated with increased gene expression. Importantly, their hypomethylated profile and enhanced gene expression were retained into DETRAINING and RETRAINING.

**Figure 4.**
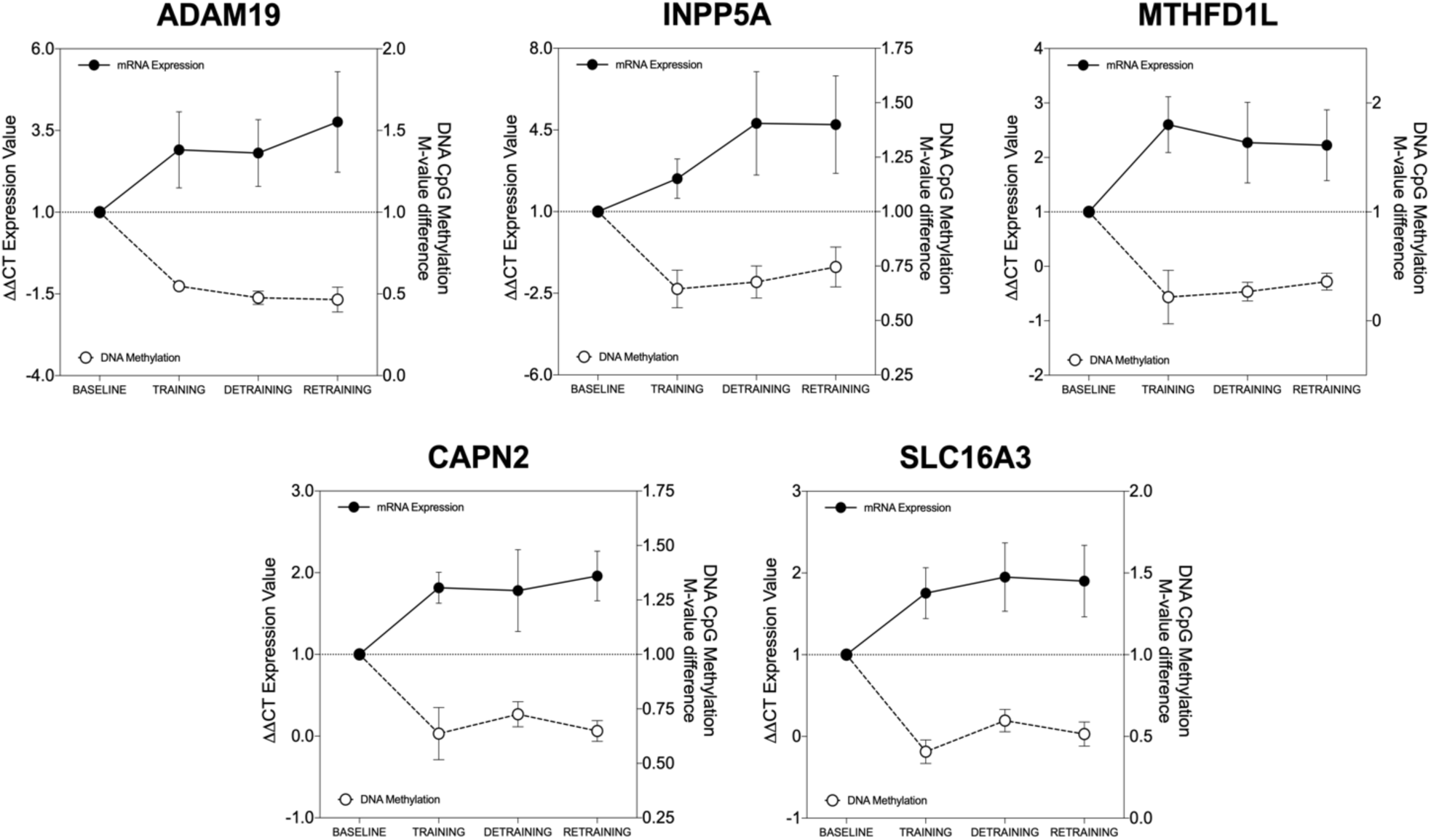
Relationship between fold changes in mRNA expression (solid black line and black dots; left y axis) and m-value difference in DNA methylation (mean m-value of all DMPs within each gene identified as having an epigenetic memory profile in either signature 1 or 2, relative to baseline) of 5 DMR gene identified (dashed black line and white dots; right y axis) across experimental conditions: ADAM19, INPP5a, MTHFD1L, CAPN2, SLC16A3. All data presented as mean ± SEM (gene expression data n = 15, methylation data n = 5).

## DISCUSSION

### Summary

In the present study we report that human skeletal muscle possesses an epigenetic memory of high intensity interval training. Eight weeks of high intensity interval training evoked a predominance of hypomethylation of thousands (total of 3,190) of CpG sites that were retained during 3 months of detraining in which exercise training was completely ceased. Interestingly, hypomethylation was retained also after a later period of retraining. Importantly, several genes could be identified as epigenetic memory genes as they retained enriched hypomethylation in genic regions and their expression continued to be upregulated even after 3 months of complete exercise cessation. Despite this, no signs of memory were observed in the positive and significant adaptations to training for the main respiratory, cardiovascular, and metabolic variables (e.g. V̇O_2max_, V̇_E_, RER, HR, [La]_b_, RPE).

### Differential DNA methylation after high intensity training, detraining and retraining

Consistently with our hypothesis, the high intensity intermittent nature of training induced epigenetic modifications at the DNA level. The total number of differentially methylated CpG sites increased compared to baseline after 2-months of training (21,605 DMPs), did not go back to baseline after 3 months detraining (3,854 DMPs) and increased again, although to a lower extent, after 2 months retraining (10,396 DMPs) (**Fig. 3A**). The training interventions also affected the status of DNA methylation, which appeared to gradually shift towards a predominant hypomethylated profile over time (**Fig. 3B**). Indeed, whereas after training there were 67% (14,516 DMPs) hypomethylated compared with 33% (7,089 DMPs) hypermethylated DMPs, three-months detraining not only reduced the total number of DMPs, but also changed their status in favor of hypomethylation (3,349 DMPs / 87% hypomethylated vs. only 505 DMPs / 13% hypermethylated). Moreover, following retraining, not only were the number of DMPs lower (10,396 DMPs) compared to training (21,605 DMPs), but a larger proportion (92% / 9,664 DMPs) showed hypomethylation compared with training (67%). Where, only 7% of DMPs were hypermethylated after retraining compared with 32% hypermethylated after training. Interestingly, 90% of DMPs located in promoter regions were hypomethylated after retraining, compared with 64% after training (**Suppl. File S2**). These findings are consistent with previous studies using both acute and chronic resistance exercise/training (Seaborne et al., 2018a; Turner et al., 2019; Bloquiaux et al., 2022) and acute and chronic endurance exercise (Nitert et al., 2012; Rowlands et al., 2014; Stephens et al., 2018; Gorski et al., 2023a, Gorski et al., 2023b) that all demonstrated a larger number of hypomethylated DMPs than hypermethylated. Interestingly, the changes in hypermethylated DMPs followed a different pattern following resistance training (Seaborne et al., 2018a) compared with the high intensity interval training intervention adopted in this study. Resistance training increased the number of hypermethylated DMPs which remained constant throughout both detraining and retraining. High intensity interval training also increased hypermethylated DMPs, but they almost reverted back to baseline levels following detraining and did not increase much following retraining. Therefore, it could be hypothesized that the two modes of exercise differ as regard the impact these DNA methylation profiles have on gene down-regulation compared with up-regulation (discussed below).

### Epigenetic memory signatures

Site specific CpG methylation and temporal analysis identified two time-related signatures of DNA methylation patterns supporting the evidence of epigenetic memory, namely, the hypomethylated profile observed in signature 1 and 2 (**Fig. 3D**), whose profile, following the initial training period, this persisted at the same loci even after 3 months of exercise cessation (detraining). Furthermore, such hypomethylation was maintained at the same sites throughout the retraining phase. These temporal trends, identify an epigenetic memory of high intensity interval training. A similar temporal profile was previously found by Seaborne and colleagues (2018) (Seaborne et al, 2018a) in response to resistance training. In this study authors identified the presence of an epigenetic memory at the DNA methylation level from a previous training-induced hypertrophic stimulus leading to retention of hypomethylation during detraining and into retraining.

Although hypomethylation was the predominant epigenetic memory signature following high intensity interval training, it is also worth pointing out that there were genes that demonstrated retained hypermethylation after training, into the detraining and retraining periods. However, this occurred in only 469 CpG sites compared with 1,190 CpG sites in signature 1 genes that show a clearly retained hypomethylation into detraining. As well as 2,000 CpGs in signature 2 and 1,175 CpGs in signature 3 that demonstrate hypomethylation with training and retraining. This retained hypermethylation in signature 4 related to only 10% that were hypermethylated vs. 90% of loci that were hypomethylated in signature 1, 2 & 3. The latter finding further supports that retained hypomethylation was the predominant epigenetic memory signature following exposure to high intensity training when training had been encountered in the past.

### Hypomethylated epigenetic memory genes in regions associated with increased transcription

As DNA hypomethylation generally leads to enhanced gene expression due to the removal of methylation allowing improved access of the transcriptional machinery and RNA polymerase that enables transcription (and also creates permissive euchromatin) (Rountree & Selker, 1997; Lunyak et al., 2002; Fuks et al., 2003; Bogdanović & Veenstra, 2009), earlier period of training should lead to increased gene expression that is retained during detraining, potentially enabling enhanced muscle adaptations in the later retraining period. In the present study, a deeper investigation of signature 1-4 genes identified several regions located in or close to annotated genes with enriched differential methylation (also known as DMRs/differentially methylated regions) that would therefore be more likely to affect gene expression. Cross referenced with the most frequently occurring CpG modifications in pairwise comparisons of all conditions resulted in identification of 11 genes whose expression was assessed by qRT-PCR. Five of such genes showed inverse relationships in methylation and gene expression (**Fig. 4**). They were related to membrane proteins involved in cell-cell and cell-matrix interactions (*ADAM19*), membrane protein that mobilizes intracellular calcium (*INPP5a)*, synthesis of tetrahydrofolate in the mitochondrion (*MTHFD1L*), calcium-activated neutral proteases calpains (*CAPN2*) and lactic acid and pyruvate transport across plasma membranes (*SLC16A3*). Expression of such genes increased after training, remained high both after exercise had completely ceased for 3 months (detraining) and after retraining. Overall, the methylation and collective responsiveness of these genes’ expression profiles identify them as epigenetic muscle memory genes following high intensity exercise training.

### Methylation and epigenetic memory genes and pathways of high intensity interval training vs. resistance exercise

The type of adaptation in skeletal muscle is dependent on the continuum of intensity for load, number of contractions, duration and frequency of exercise (Egan & Sharples, 2023). Studies to date investigating muscle memory of exercise have focused on high load, lower contraction number over a short duration, characteristic of resistance training that evokes adaptation of muscle mass and strength. This is the first study investigating potential epigenetic modifications induced by low-medium loads, higher contraction number and durations, characteristic of HIIT and endurance training. The latter training type elicit adaptations in substrate utilization, oxidative metabolism, mitochondrial function and fatigue resistance (Egan & Sharples, 2023). At the molecular level, the mechanisms responsible for differential adaptations with varying exercise stimuli are associated with divergent molecular signals, sensors and effectors. Therefore, the transcriptional and translational output that contribute to such divergent cellular and tissue adaptations are different (Egan & Sharples, 2023). However, there could be some crossover, crosstalk and common mechanisms depending on where a given type of exercise falls along the continuum of load vs. contraction number/duration. At the epigenetic DNA methylation level, it appears that divergent cellular and tissue adaptations are reflected in the gene pathways that are differentially methylated and altered at the expression level following different exercise patterns. On the heavy load resistance end of the continuum such pathways include growth, actin cytoskeletal, matrix/focal adhesion, mechano-transduction, PI3k-Akt and TGF-beta pathways in humans (Seaborne et al., 2018a; Turner et al., 2019; Sexton et al., 2023) and the mTOR pathway in mice (VonWalden et al., 2020). At the opposite end of the continuum, aerobic exercise elicits differential methylation in insulin, AMPK, substrate oxidative/fat metabolism pathways and metabolic processes (Rawlands et al., 2014; Gorski et al., 2023a). In the middle of the exercise intensity continuum between resistance and aerobic exercise, is high intensity interval training. The latter combines heavy-to-severe loading and moderate contraction number over shorter periods that are repeated. In the present study high intensity interval training evoked differential methylation in 8 common pathways within the top 20 ranked pathways across training, detraining and retraining including: (1) axon guidance, (2) calcium signaling pathway, (3) cholinergic synapse, (4) circadian entrainment, (5) focal adhesion, (6) MAPK signaling pathway, (7) pathways in cancer, (8) Rap1 signaling pathway (**Suppl. File S4**). The 7 common pathways in the top 20 KEGG pathways enriched for differential methylation across resistance training, detraining and retraining in Seaborne and colleagues (2018a) include: (1) adrenergic signaling in, cardiomyocytes, (2) aldosterone synthesis and secretion, (3) focal adhesion, (4) glutamatergic synapse, (5) MAPK signaling pathway, (6) oxytocin signaling pathway, (7) rap1 signaling pathway. Therefore, it can be suggested that focal adhesion, MAPK and rap1 signaling are common differentially methylated pathways after both resistance and high intensity interval training, detraining and retraining. Interestingly, focal adhesion, regulating ECM and mechano-transduction genes, have been previously demonstrated to be hypomethylated and upregulated at the transcriptome level after resistance training (Turner et al., 2019). Furthermore, data for differential methylation pathway enrichment here reported also overlaps with data from acute sprint interval training. Following the latter training, differential methylation in pathways that are also associated with resistance exercise such as focal adhesion pathways and axon guidance were observed (Maasar et al., 2021). Finally, acute sprint interval exercise also showed differential methylation of pathways previously observed with endurance exercise such as insulin and AMPK pathways (Maasar et al., 2021). Interestingly, MAPK pathway seems to be differentially methylated across the whole continuum, from resistance to high intensity/sprint interval and endurance exercise (Seaborne et al., 2018a; Maasar et al., 2021; Sexton et al., 2023).

In terms of the enriched differential methylation in pathways occurring in the signature 1 memory genes (**Fig. 5D**) identified in the present study, the Wnt signaling pathway, implicated in muscle hypertrophic activation (Leal et al., 2011; Armstrong & Esser, 2005), was the most enriched pathway, that demonstrated most prominent retained hypomethylation after high intensity interval training into detraining and retraining (**Suppl. File S6**). This is similar to findings in mice after progressive weighted wheel running training and detraining (Wen et al., 2021). In this study, the authors were able to demonstrate retained DNA methylation profiles form earlier training into detraining periods and investigated this in both myonuclei and all other nuclei (termed interstitial nuclei) that are resident in skeletal muscle tissue, otherwise known as interstitial nuclei (Wen et al., 2021). Indeed, Wnt signaling was identified as demonstrating an epigenetic memory profile in mouse muscle and interestingly Wnt-related genes demonstrated retained hypomethylation within the myonuclei, compared with retained hypermethylation in promoter regions within the interstitial cell nuclei (Wen et al., 2021). Therefore, the present human data also confirms that Wnt signaling is a pathway that demonstrates an epigenetic memory after high intensity interval training across all nuclei resident in human skeletal muscle tissue and supports the evidence for retained hypomethylation observed in the myonuclei of mice.

In reanalyzing data from Seaborne et al., (2018) after resistance training, detraining and retraining, we identified 5,033 DMPs that demonstrated the same epigenetic memory profile as the 1,190 DMPs in signature 1 of the present high intensity interval training study (i.e. the “retained hypomethylation” profile) after training, detraining and retraining. Running pathway analysis on such 5,033 DMPs and overlapping it with top 20 KEGG pathways from signature 1 memory genes in the present study (**Suppl. File S6**), we identified 5 pathways which were enriched in both data sets. They included: Adherens junction, Glutamatergic synapse, Parathyroid hormone synthesis, secretion and action, Proteoglycans in cancer, Ras signaling pathway. We subsequently overlapped the individual 1,190 memory CpGs from signature 1 (**Fig. 5D**; **Suppl. File S5**) that demonstrate the most prominent retained hypomethylation after high intensity training into detraining and retraining, with all those that demonstrate the same profile (5,033 DMPs) after resistance exercise training, detraining and retraining in Seaborne et al., 2018. Despite this common overlap with the pathway enrichment of hypomethylated memory genes, only 5 CpG sites were shown to be commonly epigenetic memory genes: RTTN, SRC, DDR1 and 2 CpGs on unannotated genes. Interestingly, both SRC and DDR1 are associated with focal adhesion, mechano-transduction and ECM remodeling. The latter observation suggests that different genes could be responsible for common retained hypomethylation within similar pathways after high intensity training versus resistance training. Despite this, it is important to underline that SRC and DDR1 emerge as training type independent memory genes that warrant further investigation. Finally, Filamin B (FLNB) also indicated as memory gene after resistance exercise (Turner et al., 2019) was identified in the memory signature 1 genes in the present study suggesting that filamin B could be another training type independent memory gene. Collectively, our findings and analyses confirm that muscle memory to different exercise types is mainly manifested through similar temporal patterns of hypomethylation of DMPs and enhanced expression of memory genes. However, as expected, based on their divergent functional impact, different exercise regimens activate different memory pathways and genes. Interestingly, some memory genes appear independent from the type of exercise and might support structural and functional adaptations common to all types of exercise. Finally, albeit much less predominant, hypermethylation and decreased gene expression appears a potentially interesting memory signature to focus on in future studies.

### Limitations

In the present study, high intensity interval training composed by a combination of high intensity and sprint interval sessions was undertaken (Robach et al., 2014; Granata et al., 2016). The intervention was effective since V̇O_2max_ and W_max_ were higher than baseline after both training and retraining (**Table 2**) as previously shown (Gibala et al., 2012; Granata et al., 2016). However, there seemed to be no enhanced physiological response in the retraining, compared to the training period. Although this seems to be in accordance with previous repeated high intensity interval training intervention studies (Simoneau et al., 1987; Del Giudice et al., 2020), several possibilities may account for the observed finding. One explanation could be a ceiling effect, wherein the physiological response in individuals reaches a maximum during the initial training period, leaving little room for further enhancement after retraining. This aligns with the concept that the responsiveness to training diminishes once maximum trainability is achieved (Bouchard et al., 1986). If this is the case, it might be speculated that the memory could show through a faster adaptation during the second training period, consistently with findings from repeated resistance training (Taaffe & Marcus, 1997; Staron et al. 1991; Seaborne et al., 2018a). Unfortunately, as no measurements were taken at the retraining midpoint in this project, this remains unanswered and further studies are needed. Another potential explanation could be related to the temporal duration of the 2-month interventions. It is possible that this timeframe is insufficient for the adaptive memory processes, occurring at the epigenetic and transcriptional levels, to fully manifest at the phenotype level. Finally, given the genes that manifest as epigenetic memory genes following high intensity interval training, it would be important to investigate how this leads to changes or accumulation of protein levels or protein activity and how this relates to more specific functional outcomes, related to these memory genes such as energy influx, cellular respiratory, lactate shuttling and mitochondrial biogenesis.

Lindholm et al. (2016) employed a repeated unilateral endurance training model to explore whether memory mechanisms are associated with aerobic training in humans. Gene expression via RNA sequencing did not show evidence supporting memory at transcriptional level (Lindholm et al., 2016). However, epigenetic modifications such as DNA methylation was not investigated. However, the finding here reported that several hypomethylated genes had enhanced gene expression and retained such enhanced expression even during detraining is still not consistent with Lindholm et al (Lindholm et al., 2016). Firstly, the detraining period was 9 months compared to 3 months in the present study, perhaps suggesting that epigenetic memory of aerobic training would not occur after as long as 9 months. Secondly, the exercise paradigm used by Lindholm et al., (2016) included moderate intensity activities while DNA methylation and gene expression involved in targeted metabolic and oxidative pathways seem to be more affected by high intensity aerobic exercise (Barrès et al., 2012). Interestingly, acute sprint interval exercise at a higher intensity has also been found to be more effective in impacting DNA hypomethylation changes across the methylome (Masaar et al. 2021). Therefore, it could be hypothesized that exercise for longer durations at lower loads and intensities (Lindholm et al., 2016) compared with higher intensities at high loads (present study) may be less effective in evoking an epigenetic and transcriptional memory. Despite this, genome wide DNA methylation after training, detraining and retraining at lower vs. higher aerobic exercise training of the same duration and detraining period would be required to confirm such a hypothesis. Finally, transcriptome analysis in the present study would have also provided a better understanding of the relationship between epigenetic and whole transcriptomic adaptations.

## CONCLUSIONS

Human skeletal muscle possesses an epigenetic memory (via DNA methylation) of high intensity interval training characterized by retention of DNA methylation from earlier training into detraining and retraining.

## ACKNOWELDGMENTS

We thank the participants for joining the project and Matrix Fitness Italia for their crucial contributions, including providing access to FitActive facilities equipped by Matrix and supplying MyZone MZ3 heart rate chest bands for training monitoring.

## GRANTS

Simone Porcelli was supported by a grant from Sports Medicine Italian Federation (FMSI01092021). Adam P. Sharples is currently supported by Research Council Norway (project no: 314157).

Roberto Bottinelli is currently supported by Italian Ministry for Research (MUR, PRIN Prot. 2020EM9A8X).

## DATA AVAILABILITY

DNA methylome is deposited and is freely available via Gene Expression Omnibus (GEO) with accession GSE268211.

